# Functional temporal cell type diversity in *Drosophila* reflects natural motion statistics

**DOI:** 10.64898/2026.04.30.721922

**Authors:** Luisa Ramirez, Julijana Gjorgjieva, Marion Silies

**Affiliations:** Institute of Developmental Biology and Neurobiology, Johannes Gutenberg University Mainz, Mainz, Germany; Institute for Quantitative and Computational Biosciences (IQCB), Johannes Gutenberg University Mainz, Mainz, Germany; School of Medicine and Health, Institute for Neuroscience, Technical University of Munich, Munich, Germany; School of Life Sciences, Technical University of Munich, Freising, Germany

## Abstract

As animals move through the world, their visual input can change gradually or shift rapidly, spanning a broad range of temporal frequencies. Visual circuits therefore face the challenge of encoding inputs with an inherently multiscale temporal structure. From insects to vertebrates, peripheral visual neurons exhibit striking temporal diversity, yet the principles underlying this organization remain unclear. Here, we show that temporal diversity in the fly visual system supports efficient coding of natural motion by distributing encoding across complementary timescales. Using well-characterized fly motion-detection pathways as a model, we show that temporal filters within and across visual hierarchies occupy a low-dimensional feature space defined by filter polarity and characteristic timescale. Within an efficient-coding framework constrained by natural motion statistics, circuits that combine temporally distinct filters encode naturalistic inputs more efficiently than circuits composed of more similar filters, an advantage that largely disappears when changing the stimulus statistics. Measured filters of fly visual cell types closely match the optimal solutions predicted for natural motion, and these optima map onto both circuit architecture and function. Finally, extending beyond the motion pathway, we identify distinct correlation structures in circuits within and outside the motion pathway. Together, we identify temporal diversity in response filters as a circuit-level strategy for encoding natural motion, shaped by the multiscale statistics of the environment.

**Significance statement:** Neuronal diversity is ubiquitous across sensory systems, yet its functional role remains poorly understood. We investigate a prominent form of diversity: the wide temporal response profile of visual neurons, from transient detectors to slow integrators. Using the fly visual system, we uncover a coding function of temporal diversity. Neurons with complementary temporal properties span the multiscale structure of natural motion, achieving coding efficiency that no single neuron type can implement alone. Predictions from an efficient coding framework grounded in natural motion statistics closely match fly visual circuit organization, from cell-type response properties to anatomical wiring. Because similar temporal diversity exists in the vertebrate retina, we suggest this distributed temporal coding is a conserved design principle shaped by natural scene statistics.

## Introduction

In the natural world, animals must extract behaviorally relevant information from sensory streams that evolve over time. The statistics of the inputs an animal encounters can change abruptly, such as when an object moves past the animal, or with context and self-generated actions, such as when an animal is navigating an odor plume. Consider a fly navigating a forest (Fig. 1A): Its flight trajectory can take the animal from regimes dominated by rapid image transients, e.g., vegetation moving quickly while flying close to the ground, to regimes dominated by slower fluctuations, e.g., a tree shade drifting as the sun moves. Thus, the temporal-frequency content of visual inputs is not a fixed property of the world: it is jointly determined by scene structure and the animal’s movement [1–3]. Understanding how early visual circuits support reliable behavior therefore requires linking the statistics of dynamic natural scenes to the circuit mechanisms that shape temporal coding at the earliest stages of visual processing.

**Figure 1:**
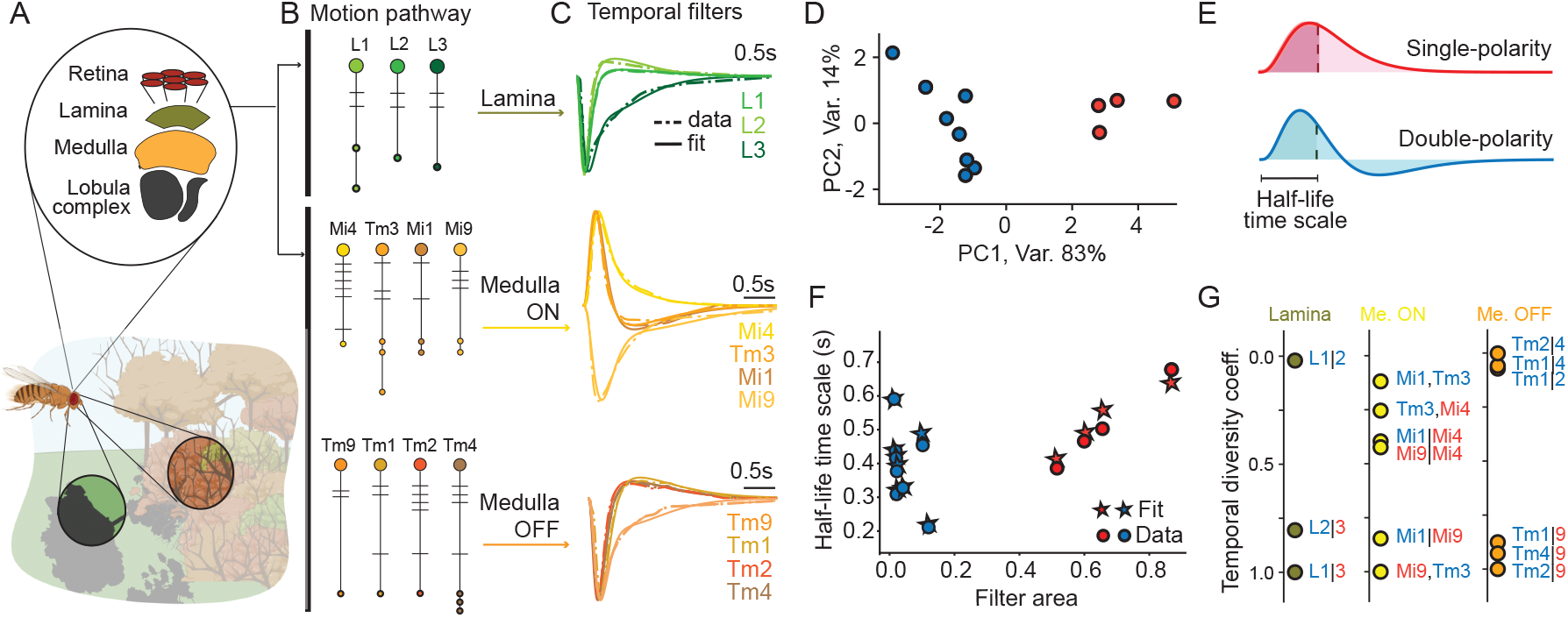
Temporal diversity in the fly optic lobe is represented by a two-dimensional feature space. **A** Schematic of a fly navigating through a natural environment, highlighting two visual patches with different temporal statistics: one containing rapidly varying motion cues, e.g. bushes, and another containing slowly varying cues, e.g. a drifting tree shade. A zoom-in shows the organization of the optic lobe neuropiles from retina to lamina, medulla, and lobula complex. **B** Schematic of the motion detection pathways considered here, showing lamina neurons and medulla neurons in the ON and OFF pathways. **C** Neuron temporal filters in the lamina and medulla. Dashed lines show the experimentally recorded filters extracted from [12], and solid lines correspond to fits to the biphasic canonical function (see Eq. 6 in Methods and Fig. S1). **D** Projection of filter data onto the PCA space. Data are color-coded by filter polarity: red for single-polarity and blue for double-polarity filters, with representative examples shown in **E**. The first two PCs reveal two main clusters characterized by filter polarity. **E** Example single-polarity (red) and double-polarity (blue) filters corresponding to the two clusters in **D**, with the half-life timescale indicated for each filter class. **F** Feature space of temporal diversity, characterized by the half-life timescale versus filter area for all fly filters, quantified by Eqs. (7) and (8). Red/blue denote single-/double-polarity filters, shown both for the experimentally derived filters (circles) and their corresponding fits (stars). **G** Temporal diversity coefficients across visual hierarchies, calculated as the normalized Euclidean distance between pairs of filters in the temporal feature space in **F** (Eq. 23). Labels show all neuron pairs, e.g. a circuit with L1 and L3 neurons is notated as L1—3. Label colors follow the polarity code in D-F.

A striking feature of early vision, from insects to mammals, is the rich diversity of temporal response properties expressed within and across the first stages of visual processing. In the vertebrate retina, temporal receptive fields range from sustained to highly transient and vary systematically across cell types, and even among neurons within the same anatomical layer [4–7]. In the mouse retina, for example, temporal diversity is already prominent at the bipolar-cell layer, where bipolar cells split the photoreceptor signal into ∼ 14 functionally distinct channels [6, 7]. This diversity extends beyond a single stage: temporal filtering is further shaped by inputs from amacrine cells, and it is then received by a diverse set of retinal ganglion cell types [4]. In insects, and in particular in the fly visual system, distinct, cell-type-specific temporal filters have been characterized within and along the visual hierarchy of the motion detection pathway (Fig. 1A-C). Pronounced temporal diversity is again evident already at the first synapses downstream of photoreceptors, in the lamina [8–10], and is then further expanded in subsequent processing layers, such as the medulla. This temporal diversity is especially well characterized in neurons of motion-detection circuits in the ON and OFF pathways (Fig. 1B,C) [8–18]. Together, these findings establish that temporal diversity is a conserved and prominent design feature of early visual systems.

Yet, the principles that generate and justify the existence of distinct functional temporal types remain poorly understood. Efficient coding theory has been successfully used to explain aspects of temporal filtering at the single-neuron level in peripheral sensory systems, where the match between response dynamics and input statistics can be formalized in analytically tractable settings [19–23]. However, visual information is processed by multi-layered circuits, with each layer combining cell types with different temporal dynamics, which are then routed into specific downstream computations. At this circuit level, it remains unclear if temporal diversity primarily reflects biophysical constraints and developmental programs, if it emerges as a byproduct of wiring and synaptic motifs, or if it represents an adaptation to natural stimulus statistics and behavioral demands. Closing this gap requires studying how combinations of distinct temporal filters perform under ecologically relevant natural motion statistics.

The pronounced temporal diversity observed at the earliest stages of visual processing suggests a functional role in information coding. Here, we explore the hypothesis that circuit pathways operating at different timescales have emerged during evolution to efficiently encode behaviorally relevant visual inputs that span a wide range of temporal scales. A key motivation is that natural motion is not well described by a single characteristic time constant: across scenes and behaviors, temporal correlations are typically broad and approximately scale-free over relevant frequency bands, consistent with power-law-like spectra [2, 22]. For such statistics, coding strategies that combine multiple timescales become natural candidates. This idea resembles other neural systems in which several adaptive processes — each well captured by an exponential with its own characteristic time constant — combine to approximate scale-free dynamics over broad temporal ranges [24–27]. In early vision, an analogous strategy could be implemented by parallel neuronal types whose temporal filters range from differentiating to integrating, collectively spanning the temporal structure of natural inputs. The fly visual system–where temporal filters are known across many identified cell types and circuit motifs can be read out from connectomes [28, 29]– therefore provides an ideal system to test how multiscale temporal diversity supports coding under natural motion statistics.

In this work, we combine data-driven modeling and theory to link natural stimulus statistics, temporal filter diversity, and circuit motifs in fly visual circuits. Characterizing the diversity of temporal filters across neuronal cell types of motion-detection pathways reveals a low-dimensional feature space, in which filters are well explained by filter polarity and neuronal response timescales. Using an analytically tractable model of dynamic natural inputs that preserves the temporal-frequency structure of natural motion, we quantify how circuits composed of temporally diverse neuron types encode natural motion under an efficient coding framework. Our analysis shows that temporal filter diversity in fly visual circuits improves the encoding of natural motion. Explicit theoretical predictions linking combinations of temporal filters to the efficient coding of natural stimuli are consistent with experimental observations. Specifically, we relate these coding predictions to the anatomical organization of motion detection circuits and to the contribution of these neuronal cell types to the function of downstream direction-selective neurons. Visual circuits not involved in motion detection appear to implement distinct coding strategies, reflected in different connectivity structures. Together, our results argue that temporal diversity in the fly visual system reflects a coding strategy tuned to natural motion statistics, in which neuronal temporal properties collectively span the multiscale temporal structure of natural motion.

### Characterization of temporal diversity in the fly visual system

Peripheral neuronal cell types in the fly visual system exhibit a wide range of temporal responses, both within and across visual hierarchies. To quantify this diversity of temporal features, we analyzed previously reported temporal filters of cell types in the fly motion detection pathway [12]. Within visual hierarchies of the achromatic pathway, temporal response diversity is first apparent in the lamina and medulla neuropiles (Fig. 1A-C), where neuronal response functions have been extensively characterized through linear temporal filters [12]. Combining these peak-normalized filters (Fig. 1C), we performed a principal component analysis (PCA) and found that the first two principal components capture 97% of the variance (Fig. 1D and Methods), indicating that the temporal diversity is well described in this low-dimensional space. This two-dimensional representation separated the filters into two main clusters corresponding to single (monophasic) versus double polarity (biphasic) temporal profiles. The first principal component (PC1) primarily distinguishes these polarity types, while the second (PC2) captures systematic variation within each polarity type (Fig. 1D). Consequently, we interpreted PC1 in terms of filter area, which increases systematically when moving from double-polarity to single-polarity filters (Fig. 1E). In contrast, PC2 varied more continuously and captured differences in the filters’ mean response time, which we related to each filter’s half-life timescale (Fig. 1D). To quantify these two features, we fitted all filters with a canonical biphasic function [30] (Methods and Figs. 1C). From these fits, we extracted filter area and half-life timescale (Fig. 1E,F and Eqs. (7)-(8)) in Methods), yielding a feature space that preserved the PCA clustering, while providing axes with clear physiological meaning. The principal axis reflects polarity, spanning responses from more differentiating (double polarity) to more integrating (single polarity), while the orthogonal axis captures differences in characteristic time scales and accounts for most of the variability within each polarity type (Fig. 1F).

Next, we examined the distribution of temporal diversity within visual hierarchies. We defined the temporal diversity coefficient of a given circuit as the normalized Euclidean distance between filters in the temporal feature space (Fig. 1G and Eq. 23). For two-neuron circuits, we calculated this coefficient within each hierarchical level. In the lamina, temporal diversity is highest for the pairs {L1, L3} and {L2, L3}, whereas {L1, L2} exhibits substantially lower diversity. In the down-stream medulla, ON and OFF circuits span a comparable overall range of temporal diversity values. However, two-neuron circuits in the ON pathway distribute more uniformly across this range, while OFF-pathway pairs more prominently cover the extremes with either low or high diversity (Fig. 1G). Altogether, our analysis reveals that the temporal diversity in the fly visual system is organized primarily along two axes: response polarity and characteristic timescale, providing an accurate description of temporal responses within and across two peripheral visual hierarchies.

### Dynamic natural inputs exhibit multiple relevant temporal scales

The pronounced temporal diversity observed early in visual processing suggests a functional role in information coding. We propose that circuit pathways operating at different timescales may have emerged due to evolutionary pressure to efficiently encode behaviorally relevant visual inputs that span a wide range of temporal scales. This idea parallels other neural systems in which multiple adaptive processes – each well approximated by an exponential with its own characteristic time constant – combine to approximate scale-free dynamics over extended temporal ranges [24–27]. Investigating whether early visual pathways implement a similar strategy requires linking the statistics of natural motion with the temporal properties that determine encoding in fly visual circuits. We thus first characterized the statistical structure of natural motion using an analytical framework that goes beyond individual examples (e.g., specific video recordings) and yields general insights. Natural motion, as seen from a fixed viewpoint, produces luminance and contrast fluctuations with power spectral densities that follow power-law statistics (Fig. 2A,B) [2, 21, 22]. Such correlation structure means that informative fluctuations are distributed across a wide range of temporal scales. Across natural scenes, ranging from waterfalls to insect swarms, spectral densities yield different power-law distributions with exponents between k = −2 and k = −1 (black curves in Fig. 2B) [2]. To relate the temporal diversity in fly visual circuits to the temporal-frequency structure of natural motion, we focused on a simplified, analytical description that preserves this structure at the level of its power-spectrum, rather than the full spatiotemporal content of natural movies. Specifically, we generated signals through a mixture of low-pass exponential filters with temporally white input, that recovered the observed spectral power-law within relevant two decade frequency ranges reported in the literature (Fig. 2C, Methods). We constrained the model to ecologically relevant temporal frequencies based on measurements from natural movies (Fig. 2B) [2]. To attenuate frequencies outside the experimentally reported range without affecting the power-law within the two-decade frequency range observed, we added a band-pass filtering stage to the low-pass mixture (Fig. 2C and Methods). Varying the number of exponential stages in the mixture from one to three allowed us to generate a set of power-laws with exponents ranging from −2 to −1. For each mixture, we fitted the low-pass filter parameters (amplitudes and timescales) to match the target spectra. In practice, a two-filter mixture was sufficient to reproduce the observed power-law behavior over the two decade temporal frequency range observed in data (Fig. 2D). We settled on a minimal model where we fitted only one of the parameters of the mixture model, e.g., the timescale of the second low-pass filter, reproducing a set of power laws that reflects natural dynamic scene statistics (Fig. 2E).

**Figure 2:**
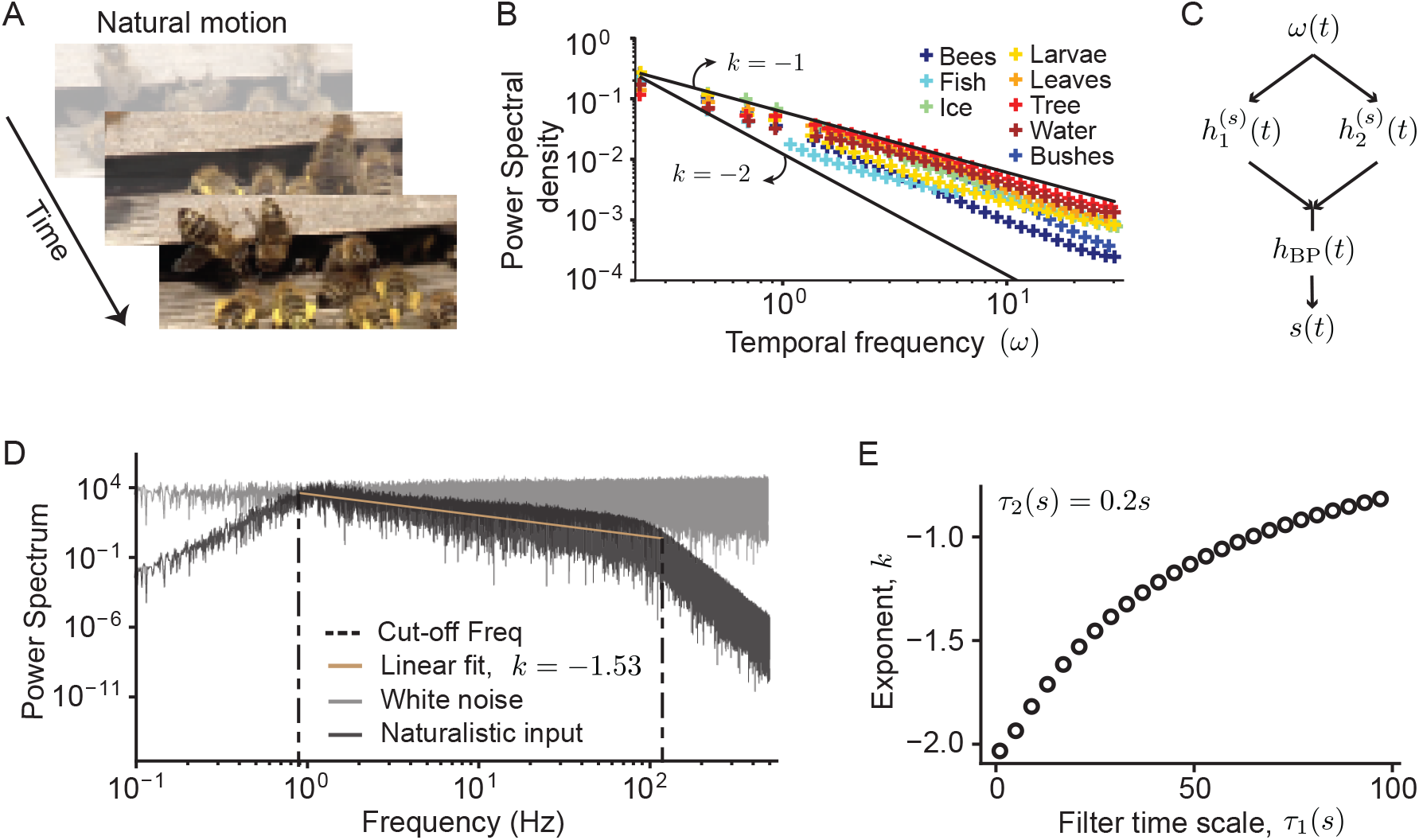
Characterization of dynamic naturalistic stimuli. **A** Frames at different time points of a natural video of moving bees [31]. **B** Power spectral density versus temporal frequency for different natural movies; colored markers reproduce data extracted from [2]. Black curves show power law distributions, *∝* ω^*k*^, with exponents k = −2 and k = −1. Data points fall within this range [2, 22]. **C** Two-stage mixture of low pass linear filters, 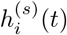, that generate a naturalistic stimuli, s(t), from white noise, w(t). A band-pass filter, h_*BP*_ (t), restricts the signal to frequencies reported in the literature (Eq. 4). **D** Power spectrum of a white noise (gray) and a naturalistic (black) signal generated by the mixture of low-pass filters in C. The mixture transforms the input (gray) into an output (black) with an approximate power-law with exponent K ∼ − 1.5 within the frequency range from data, temporal frequencies outside this range are attenuated by the band-pass filter. **E** Power law exponents produced by the mixture of two filters. Varying a single timescale, τ_1_(s), while keeping the timescale of the second filter as well as the amplitudes fixed (τ_1_ = 0.1, α_1_ = 100 and α_2_ = 0.1) captures the range of exponents that characterize natural motion.

### Temporal diversity in lamina and medulla networks improves encoding of natural motion statistics

To test whether the temporal-frequency structure of natural motion has shaped the emergence and organization of temporal diversity in the fly optic lobe, we next investigated the coding performance of circuits with different degrees of temporal diversity in the most peripheral lamina and medulla neuropiles (Fig. 1F,G). Within each visual layer, we modeled circuits as sets of neuronal types with distinct temporal filters driven by stimuli with naturalistic power-law power spectra (Eq. 5, Fig. 2B). A linear encoder, optimized for stimulus reconstruction (Eq. 21 in methods), integrated these neuronal responses to generate the network output (Eq. 9, Fig. 3A). We assessed the coding reliability of the circuits through the signal-to-noise ratio (SNR), calculated via the mean square error between the input and the reconstructed stimulus (Methods and Fig. 3A). This performance metric provides an analytically tractable and comprehensive measure of how well the circuit preserves stimulus information across the relevant frequency band [20, 32–34].

**Figure 3:**
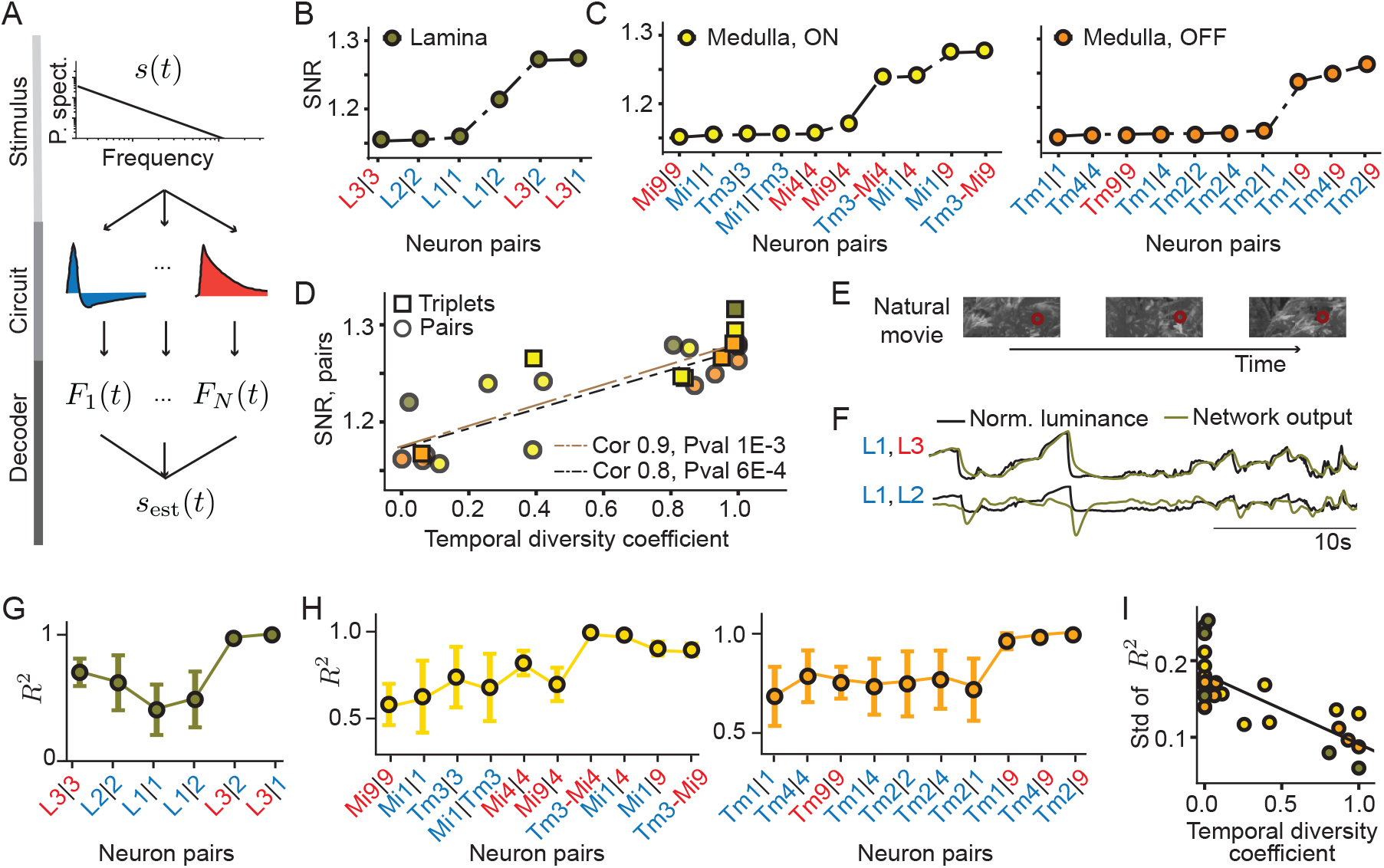
Temporal diversity in the fly peripheral optic lobe enhances the encoding of naturalistic stimuli. **A** Network coding schematic. The naturalistic stimulus, s(t), is filtered by the linear filter of each neuron and then read out by an optimized linear decoder, **F**(t), that generates the network output, S_est_(t). **B-E** Analytical results for lamina and medulla networks (Methods). **B** Signal to noise ratio (SNR) for lamina networks of size N = 2; cell type names are color coded by filter polarity type as in Fig. 1D-G. Neuron filters were fit with function Eq. (6) used in Fig. 2, and the optimization was done over the linear decoder, **F**(t) (Methods). **C** Same as in B for ON and OFF medulla networks. **D** SNR versus temporal diversity coefficient of Eq. 23 for non-redundant network combinations in the lamina (green), the ON-medulla (yellow) and the OFF-medulla (orange). Circle (square) markers correspond to circuits composed of neuron pairs (triplets) (see also Fig. S2). **E-I** Simulations using luminance traces extracted from natural movies (Methods). **E** Example scenes from one video. Luminance traces were extracted for specific pixels after applying a smoothing spatial Gaussian filter of σ = 6 pixels. **F** Luminance stimulus (black) and network response traces (green) for two network examples in the lamina with predicted high and low SNR in C. **G** Encoding performance for each lamina network, quantified by the R^2^ of a linear encoding model (temporal lag window of 1s). Networks are sorted as in C. Error bars extracted from simulations across 11 natural videos [2,31]. **H** Same as in G for ON-medulla (left) and OFF-medulla (right) networks. **I** Std of R^2^ versus temporal diversity coefficient across all networks (colors as in B-C). Lamina and medulla networks with large temporal diversity exhibit a high encoding performance with higher reliability (small std) across natural motion scenes. (see also Fig. S2 for circuits with neuron triplets). All SNR calculations correspond to a power-law stimulus statistics with k = −1.57 (Fig. 2). A systematic study over all different exponents described in Fig. 2 leads to similar results and is summarized in Fig. S2E

The lamina contains the double-polarity neuronal types L1 and L2, and the single-polarity neuron type L3, all receiving direct input from photoreceptors [12, 35]. We first considered lamina circuits of varying size in our efficient coding framework. In two-neuron circuits, we evaluated all possible pairs drawn from {L1, L2, L3}. We found that all lamina circuits with high temporal diversity, e.g., those combining double and single polarity filters (Fig. 1G), achieved high SNR, as in networks where L3 was combined with L1 or L2. In contrast, circuits with low temporal diversity, such as pairs with identical(L1, L1}) or similar ({L1, L2} temporal filters, exhibited lower SNR (Fig. 3B). Interestingly, combining single- and double-polarity filters consistently outperformed combinations of filters of the same polarity type, even when those differed by their half-life timescale (Fig. 3B and Fig. 1F).

We observed the same pattern in both the ON and OFF pathways of the medulla (Fig. 3C): circuits with the same polarity filters exhibited lower SNR than circuits with different polarity filters (Fig. 3B,C). To quantify the general dependence of SNR on temporal diversity, we computed a temporal diversity coefficient for each circuit (Fig. 1G and Eq. 23). Across all medulla and lamina circuits with non-zero diversity, SNR increased linearly with the temporal diversity coefficient (Fig. 3D). This linear relationship remained evident in both ON and OFF medulla pathways when extending to larger circuits composed of three neurons (Fig. 3D).

Together, these results show that circuits composed of neurons with diverse temporal filters (Fig. 1G) enhance the encoding of natural motion statistics across hierarchies and pathways of the fly visual system. Specifically, our findings argues that fly temporal filters may have evolved into integrators (single-polarity filters) and differentiators (double-polarity filters) to broaden the effective frequency range of circuits and support reliable coding of natural motion across a broad range of temporal scales [24–27].

We next asked whether our theoretical results for the encoding of a stimulus model, that matches natural motion only through its power-spectral statistics, generalizes to real motion sequences from natural movies. To test this, we simulated the responses of all two-neuron lamina and medulla circuits (Fig. 3B,C) to luminance traces extracted from natural videos spanning various ecological scenes, such as the movement of the leaves from a tree on a windy day or the movement of bees on a hive (Methods; Fig. 3E,F). For a given luminance trace, the lamina circuit with the highest SNR, {L1,L3}, exhibited a more accurate reconstruction of the response compared to a circuit with lower SNR, {L1, L2} (Fig. 3F). To quantify coding reliability across circuits and stimuli, we measured the match between the encoded stimulus trace and the true stimulus by the R^2^ of a linear encoding model. Furthermore, we evaluated each circuit on a set of 11 different stimuli extracted from different videos and computed the mean R^2^ and its corresponding standard deviation (std) across stimuli. Overall, lamina and medulla circuits with high SNR reliably encoded natural dynamic stimuli, which generalized robustly across different videos (Fig. 3G,H). In contrast, low-SNR circuits achieved lower R^2^ with larger variability between stimuli, likely reflecting the specific temporal frequency content of each individual video and suggesting reduced robustness to stimulus variation (Fig. 3I). Similar results were observed when studying circuits composed of three neurons.

Together, these results show that temporal diversity across circuits in the visual hierarchy enhances the coding of natural motion in the fly visual system.

### Temporal properties of fly visual cell types are matched to natural stimulus statistics

Given that temporal diversity improves the encoding of natural motion, we next asked whether the statistics of dynamic natural stimuli could have driven the emergence and organization of such diversity. We approached this question from two angles. First, we tested whether the encoding benefits of circuits with temporally diverse filters remains when changing the stimulus statistics. Second, we asked whether the temporal diversity measured in the fly visual system is close to optimal for encoding natural motion.

Addressing these questions required expanding the space of possible temporal filters beyond the set measured in the fly. We therefore generalized neuronal temporal filters within the same low-dimensional feature space that captures temporal diversity in the optic lobe (Fig. 4A), excluding biologically implausible solutions with unrealistically fast or slow response dynamics. By filling regions of the feature space not occupied by the experimentally measured fly filters, we generated a new set of temporal filters (Fig. 4A). Consistent with the low-dimensional representation of the experimental data, we preserved the categorization between single- and double-polarity filters along the area axis, such that double-polarity filter areas ranged from 0 to 0.4 and single-polarity filter areas ranged from 0.4 to 1 (blue and red lattices in Fig. 4A). This expanded filter set comprised 97 double-polarity filters, compared to 8 measured in the fly, and 94 single-polarity filters, compared to 4 measured in the fly (Fig. 4A, Methods).

**Figure 4:**
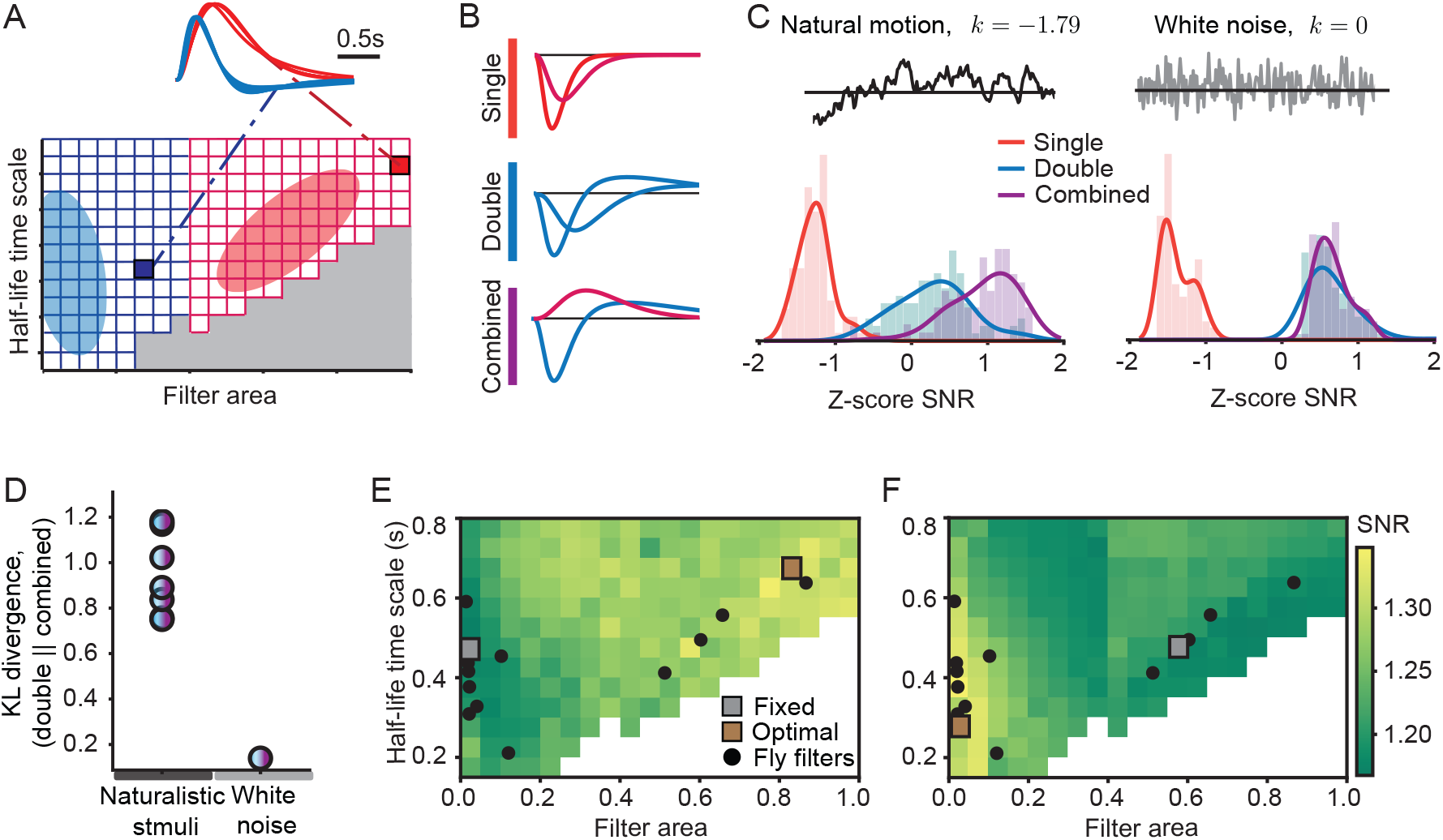
Temporal diversity in the fly visual system matches efficient coding predictions for natural motion statistics. **A** Generalized temporal filter space (step 0.05 in filter area and half-life timescale) spanning fly-like filters. Filters are generated by fitting Eq. 6 (example traces). Single-polarity filter region (red): 94 filters with area in [0.4, 1]. Double-polarity filter region (blue): 101 filters with area in [0.05, 0.4). Gray: forbidden region (large area incompatible with very short half-life). **B** Two-neuron circuit classes: single-polarity, double-polarity and combined-polarity (with one of each single and double polarity filters). **C** z-score SNR distributions for the three circuit classes (see B under naturalistic power-law input (k = −1.79) and white noise (k = 0). Top: example stimuli. Bottom: histograms and corresponding kernel density estimation (KDE) of the z-scored SNR across all filter combinations within the generalized feature space. **D** Kullback-Leibler (KL) divergence between double- and combined-polarity SNR KDEs across stimulus statistics. Left column: naturalistic power-laws with exponents in [−2, −1]. **E-F** SNR landscape for two-neuron circuits with one filter fixed (gray marker) and the partner varied across the feature space (color=SNR). The SNR-maximizing partner (brown marker) is typically of opposite polarity. **E** Fixed: double-polarity filter; optimal partner: single-polarity filter. **F** Fixed: single-polarity filter; optimal partner: double-polarity filter. Black dots: measured fly filters.

Building on the polarity-based organization of fly temporal filters, we considered two-neuron circuits (as in Fig. 3B-C) and grouped them into three main categories: I. single-polarity circuits in which both neuronal types have single-polarity filters, II. double-polarity circuits in which both neuronal types have double-polarity filters, and III. combined-polarity circuits which pair a single-with a double-polarity filter (Fig. 4B). For example, the lamina circuit {L1, L2} belongs to the double-polarity category, while {L1, L3} is a combined-polarity circuit.

To probe how temporal diversity depends on stimulus statistics, we calculated the SNR of all circuits for two main types of stimuli; natural motion constrained by the power spectral statistics and white noise (Fig. 4C and Methods). For natural motion, combined-polarity circuits achieved the highest SNR, followed by double-polarity circuits, with single-polarity circuits performing the worst (Fig. 4C, example power-law with k = −1.79). Strikingly, the higher SNR, indicating better coding performance, was specific for stimuli constrained by power-law spectral statistics; under white-noise stimulation, double- and combined-polarity circuits performed comparably (Methods).

We generalized these results by varying the power-law exponent to capture different ecological contexts (Fig. 2D,E), and calculated the corresponding SNR distributions as in Fig. 4A-C. Across all tested power-law exponents, the separation between the SNR distributions of double- and combined-polarity circuits remained robust. Quantifying this separation via the Kullback-Leibler (KL) divergence confirmed that the encoding benefits of circuits with temporally diverse filters are specifically enhanced under naturalistic, power-law stimulus statistics (Fig. 4D). The similar SNR obtained from double- and combined-polarity circuits for white noise input statistics suggests that this diversity might have evolved as a solution for the encoding of ecologically relevant motion.

Next, to test whether fly temporal filters are matched to natural input statistics, we visualized optimal encoding strategies across all two-neuron circuit combinations in the extended feature space. Specifically, for each filter in the feature space, we identified the partner filter that maximized circuit SNR. To illustrate this, we fixed a reference filter (Figure 4E, gray square) and computed the SNR of the resulting two-neuron circuit as the partner filter varied across half-life timescale and filter area (Figure 4E). Within this feature space, we identified explicitly the SNR-maximizing partner: e.g., a double-polarity reference filter (Figure 4E, gray square) was optimally paired with a single-polarity partner (Figure 4E, orange square). Both the reference filter and its optimal partner lay close to filters measured in the fly (black dots, Figure 4E). Conversely, starting from a single-polarity filter, its optimal partner was typically a double-polarity filter (Fig. 4F). Across most of the feature space, the optimal solutions selected temporally diverse pairings of single- and double-polarity filters matching observations in the fly visual system, suggesting that fly-like diversity yields near–optimal encoding under natural motion statistics.

Overall, our results support the view that the fly visual system implements an efficient strategy for the encoding of natural motion by splitting temporal processing into different channels with carefully tuned characteristic scales. In this framework, differentiating (double polarity) neuronal types and integrating (single polarity) neuronal types together cover the multiscale temporal structure of dynamic natural inputs (Fig. 4E,F and Fig. 3E). This strategy provides clear encoding benefits specific to naturalistic, power-law stimulus statistics. These benefits are significantly reduced under motion statistics not tuned to naturalistic scenes, e.g. white noise, where the observed neuronal temporal diversity would not have provided key evolutionary advantages for the encoding of motion.

### Coding efficiency is consistent with the columnar organization of medulla inputs onto direction-selective dendrites

Beyond temporal diversity, motion detection circuits in the fly visual system rely on a systematic spatial organization of medulla inputs onto the dendrites of postsynaptic direction-selective neurons [12, 37]. Having shown that the temporal properties of cell types in the fly visual system are well matched to the statistics of natural motion, we next asked how these same natural motion statistics have constrained circuit organization in the motion detection pathway. The spatial arrangement of the connectivity from medulla neurons onto T4/T5 dendrites has been described in detail (Shimomiya 2019 eLife). Both T4 and T5 dendrites receive inputs from medulla neurons spanning three neighboring columns (Fig. 5A). In the ON pathway, a T4 dendrite receives Mi4 input towards the base of its dendrite, Mi1 and Tm3 at the center, and Mi9 synapses towards the distal tip. In the OFF pathway, a T5 dendrites receives CT1 input towards its base, Tm1, Tm2 and Tm4 synapses at the center, and Tm9 synapses towards the distal tip (Fig. 5A).

**Figure 5:**
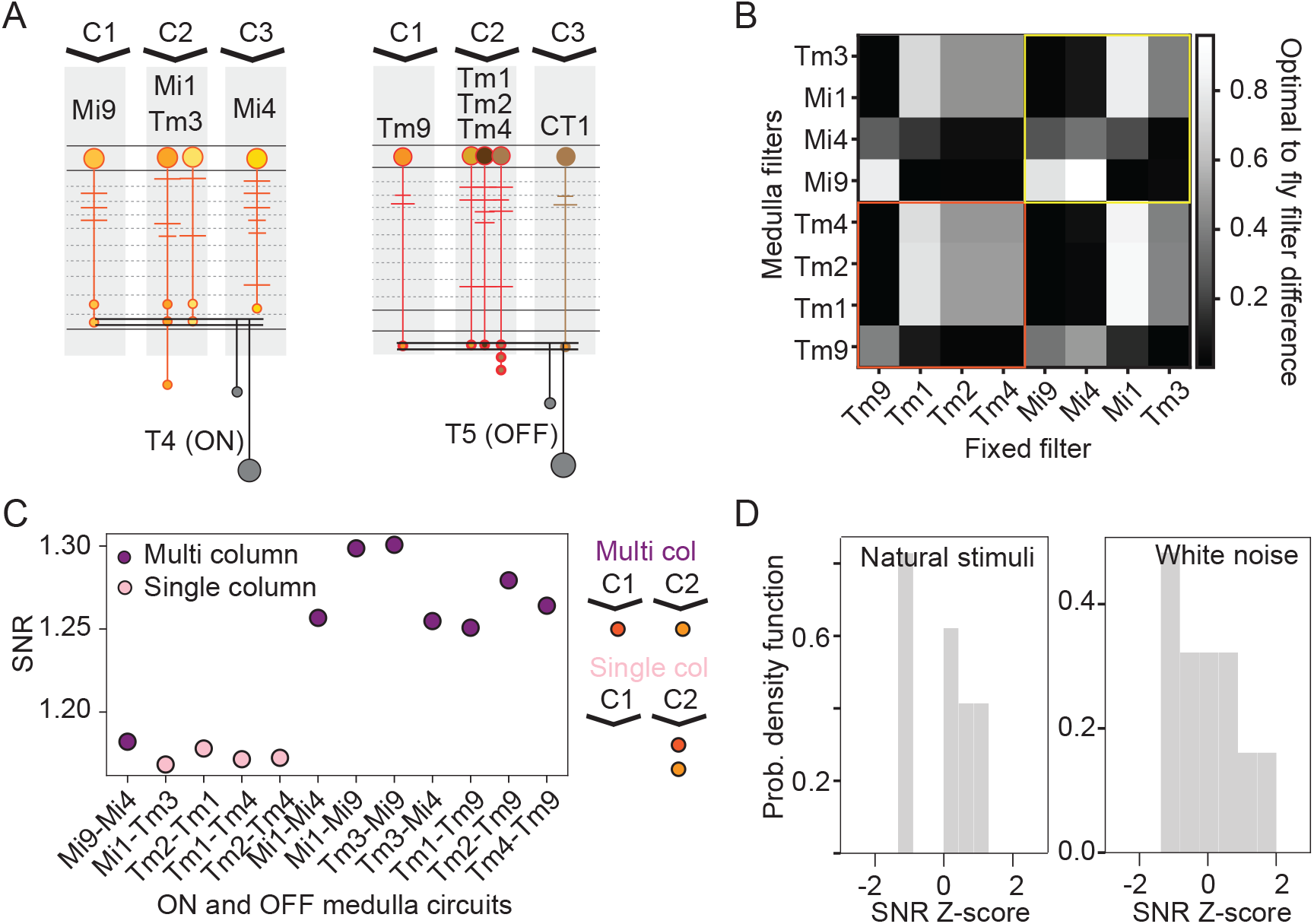
Temporal diversity in the motion detection pathway is matched to natural motion statistics. **A** Schematic of the motion detection circuit in the ON (left) and OFF (right) pathways. T4/T5 receive input across at least three columns (C1, C2, C3) and each column contains at least one columnar input to the downstream direction-selective neuron. From base to tip, these Mi4, Tm3/Mi1, Mi9 for the T4 dendrite and CT1, Tm1/Tm2/Tm4 and Tm9, for T5. The temporal filter of CT1 has not been experimentally characterized, but has been approximated by the filter of its main presynaptic partner Tm9 [36]. **B** Distance map comparing predicted optimal partner filters to measured fly filters for all medulla neurons. For each fixed medulla neuron filter (x-axis), we first computed the SNR-maximizing partner filter from the generalized feature space (as in Fig. 4E,F), then quantified its similarity to every measured medulla filter (y-axis) using a normalized square-error distance in the feature space (0 = most similar; 1 = most dissimilar; normalization by the maximal squared error over all generalized fly filter pairs).**C** SNR for all combinations of medulla neuron pairs presynaptic to T4 (ON) or T5 (OFF), sorted by their circuit type: single-column pairs, if the presynaptic neurons belong to the same column (light pink), or multi-column pairs, if the presynaptic neurons belong to different columns (purple). **D** Histogram showing the SNR distribution for all pair circuits in the ON and OFF pathway under natural stimulus statistics (left) or white noise stimulus statistics (right).

To relate this wiring motif to efficient coding under natural motion, we compared optimal coding circuits to the fly motion detection circuit organization. As above (Fig. 4E,F), we fixed the filter for each medulla neuron and identified the partner filter which maximized SNR. Within each pathway, the fly filter closest to the predicted optimal partner typically belonged to a pair in which one neuron targets the dendritic center and the other targets an adjacent position (base or tip) (Fig. 5B). For example, the predicted optimal partners of the OFF medulla neuron Tm9 (input at the dendrite tip) matched the temporal properties of the OFF medulla neurons Tm4/Tm2/Tm1 (all inputs at the dendrite center). Comparable predictions were also predicted for other pairs in the ON pathway, such as {Mi9, Mi1}. In contrast, pairs that both target the center within the same column (e.g., {Tm1, Tm4}) yielded poor SNR.

As before, although the preference for opposite-polarity partners was evident within both the ON (T4) and OFF (T5) pathways, our distance-based metric did not cleanly recover the ON/OFF circuit partition. In some cases, cross-pathway matches were as close as, or closer than, the biologically paired partners (e.g., Mi9 matched well to Tm1/Tm2/Tm4 and to Mi1/Tm3). Altogether, our results suggest that the columnar organization of motion detection circuits within the ON and OFF pathways also matches the statistics of natural motion.

To make this link explicit, we calculated the SNR for all single-column and multi-column circuits within each pathway, and color-coded circuits by whether they spanned one column or multiple columns (Fig. 5C). All adjacent multi-column circuits exhibited higher SNR than single-column circuits. The only multi-column circuit with a low SNR contained neurons more than two columns apart, {Mi9, Mi4}. Thus, temporally diverse filter combinations are organized in an adjacent multi-columnar manner, matching the known motion-detection circuit organization [37].

Finally, we asked whether this relationship between SNR and columnar organization is specific for natural motion statistics. We thus compared circuits’ SNR under natural power-law statistics versus white noise motion statistics. Under natural power-law statistics, SNR exhibited a bimodal distribution (Fig.5D), consistent with the strict separation between low-diversity single-column pairs and high-diversity multi-column pairs (Fig.5A-C). Strikingly, this two-mode structure disappeared under white noise motion statistics (Fig.5D), indicating that when motion cues lack natural temporal structure, remaining temporal diversity does not yield further coding benefits. Altogether, these results show that temporally diverse filter combinations preferentially map onto multi-column wiring motifs, linking efficient coding under natural motion statistics to the columnar organization of presynaptic inputs onto direction selective neurons.

### Temporal diversity and functional synergy exhibit different relationships in circuits within and outside the motion detection pathway

To connect our coding predictions to downstream function, we tested whether circuit SNR correlated with the effects of perturbing temporal diversity in medulla circuits. To this end, we analyzed experimental results from the OFF pathway, where the same medulla neuron types studied here were functionally assessed through acute genetic silencing [15]. Specifically, Tm1, Tm2, Tm4, and Tm9 were silenced either individually or in pairs, and visual responses were recorded electrophysiologically from neurons directly downstream of T4/T5 (the lobula plate tangential cells, LPTCs) (Fig. 6A). We quantified the effect of silencing as the LPTC peak response normalized by the control response (without silencing) (Fig. 6B). This provides a measurement of how partial medulla circuits contribute to visually evoked motion responses of downstream LPTC neurons. For example, silencing Tm9 lead to a stronger reduction in LPTC response than e.g., Tm1. We next computed the SNR of each partially silenced circuit by applying the equivalent silencing in our model, i.e., removing silenced neurons from the decoder. Notably, SNR of the remaining circuit exhibited a significant positive correlation with the strength of LPTC response across all silencing conditions (Fig. 6C), showing that downstream neurons are more sensitive to perturbations that significantly decrease the temporal diversity of the pathway than perturbations in which temporal diversity is less affected.

**Figure 6:**
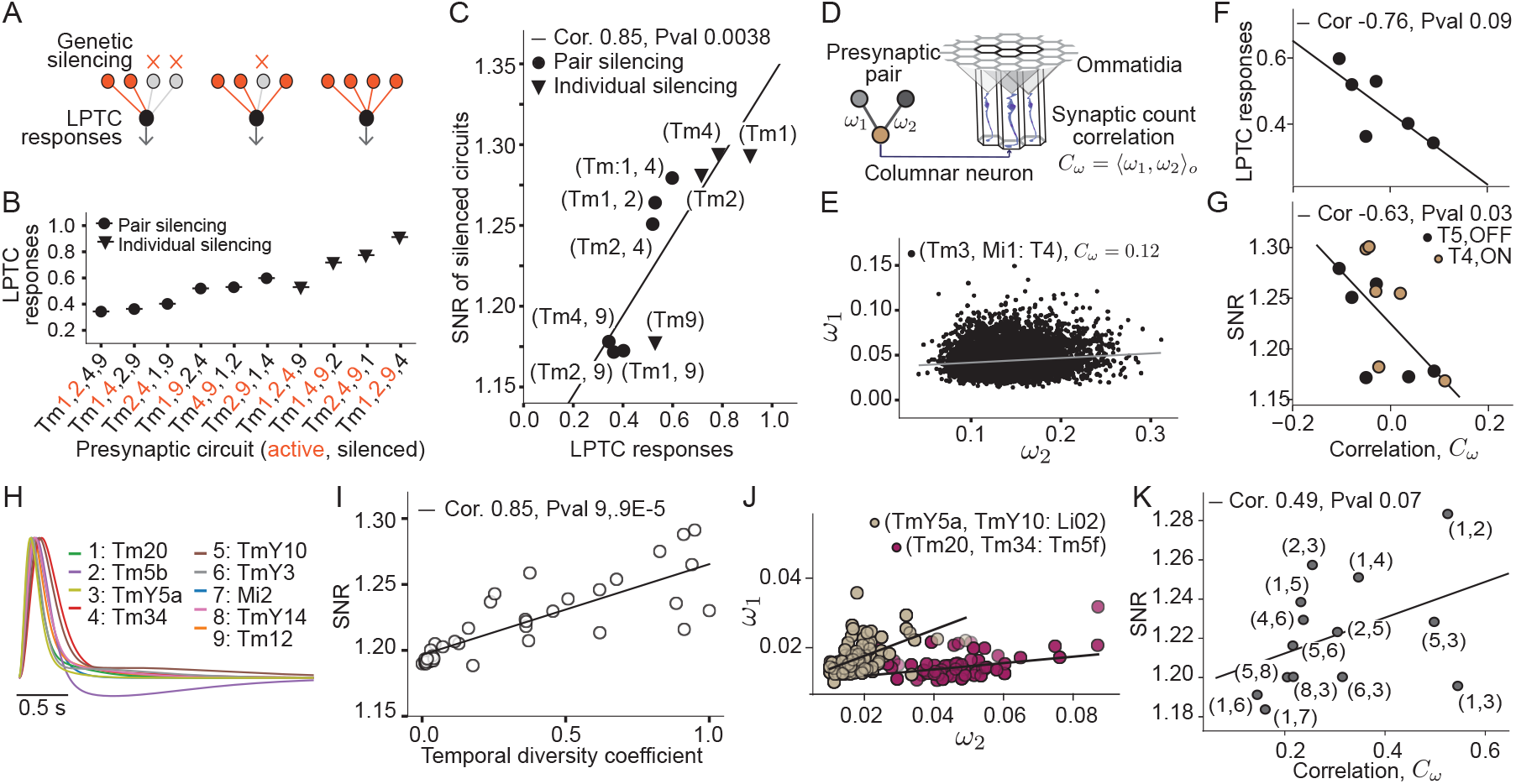
Connectivity correlations act as an intermediate measure to link temporal diversity and circuit function in the medulla neuropile. **A** Experimental design to measure the effects of perturbed medulla inputs in the motion detection OFF pathway. LPTC neuronal activity, downstream of T5 neurons, is recorded in response to the genetic silencing of either single or pairs of medulla neurons [15]. **B** Peak response of LPTC neurons under the partial silencing of the four medulla inputs in the motion detection pathway. Responses are normalized by the peak response without silenced medulla neurons. Error bars as in the experimental data extracted from [15]. Orange (black) labels correspond to the silenced (unperturbed) neurons, and circle (triangle) markers correspond to circuits with two (one) silenced neurons. **C** SNR of the silenced circuit as a function of the downstream LPTC response intensity. Markers as in B. **D** Correlation measure of synaptic counts for visual circuits in the connectome. For each circuit, we calculate the correlation between synaptic counts, ω_1_ and ω_2_, across ommatidia. **E** Synaptic counts for one circuit of the motion direction pathway across columns, with the presynaptic neurons Tm3 and Mi1 and the postsynaptic neuron T4. ω_1_ (ω_2_) corresponds to the synaptic count between the medulla neuron Tm3 (Mi1) and the postsynaptic neuron T4. Each point corresponds to a different columnar circuit. Measured correlation C_*ω*_ = 0.12 with P-val = 0.003. **F** LPTC intensity response as a function of the correlation, C_*ω*_, for all circuits composed of two neurons (circles in panel C) in the OFF motion detection pathway. **G** SNR as a function of the correlation, C_*ω*_, for all circuits composed of two neurons in the ON and OFF motion detection pathways. **H** Fit of temporal filters of medulla neurons outside the motion detection pathway. Data extracted from [12]. **I** SNR as a function of the temporal diversity coefficient (as in Fig. 3D) for circuits of size N = 2 with the new medulla neurons in H. Networks of size N = 3 also exhibit a positive correlation (see Sup. Fig 6 Update figure). **J** Same as in E for two circuits outside the motion detection pathway. Postsynaptic partners are chosen based on the ranked synaptic count and statistical significance (see Methods). **K** SNR as a function of the circuit correlation C_*ω*_, for all circuits outside the motion detection pathway. Labels as in H.

We next asked whether the functional outcomes of partially perturbed circuits are also reflected in circuit connectivity. Specifically, we tested whether partial circuits with stronger downstream effects exhibited more structured connectivity than partial circuits with weaker effects. We calculated the correlation between synaptic counts of pairs of medulla neurons presynaptic to T5 across columns (Fig. 6D), allowing us to characterize different connectivity structures, e.g. strongly correlated versus independent (or anti-correlated) connectivity patterns (Fig. 6D. For example, for the circuit where Tm3 and Mi1 connect to the postsynaptic neuron type T4 (referred to as Tm3, Mi1: T4), we extracted their relative synaptic counts across ∼ 800 medulla columns and quantified the resulting connectivity correlation (Fig. 6E. In the OFF pathway, the partial circuits that elicited the strongest reductions in LPTC responses exhibited negative correlations, whereas circuits that elicited a weaker reduction were positively correlated (Fig. 6F). Consistent with this, SNR decreased with increasing connectivity correlation in both ON and OFF pathways (Fig. 6G and Fig. 6C,F). Together, these results link coding and function in the motion pathway and show that temporally diverse circuits have a stronger impact on downstream motion responses than temporally less diverse circuits. When examining connectivity features, we found circuits with either positive or negative correlations between their synaptic counts (Fig. 6D).

Finally, to explore whether medulla circuits outside the motion detection pathway exhibit similar circuit properties, we explored a new broader set of medulla neurons within the same visual hierarchy with recently measured temporal filters [12] (Fig. 6H and methods). Because the functional roles of these neurons are not yet known, we considered all pairwise combinations and computed the SNR and the temporal diversity coefficient, as before (Fig. 3). As in the motion detection pathway, circuits with high temporal diversity achieved a higher SNR for natural motion statistics compared to less diverse circuits. We next assessed circuit connectivity organization by first identifying common postsynaptic partners for each presynaptic pair, analogous to the T4 and T5 neurons in the motion pathway. For the most significantly connected postsynaptic neurons, we measured the correlation between the presynaptic synaptic counts across neurons of the same type (Fig. 6J,K), as we did in the motion detection pathway (Fig. 6E). Medulla circuits outside the motion detection pathway exhibited only positive connectivity correlations. Furthermore, circuits with high SNR (and more temporally diverse filters) exhibited the strongest positive correlations (Fig. 6K). These strong correlations indicate that the synaptic strength of these medulla neurons is co-regulated, suggesting a potential synergistic role for shared downstream neurons in non-motion pathways. Taken together, these findings show that temporal diversity shapes connectivity structure in medulla circuits, also revealing wiring differences between motion and non-motion pathways.

## Discussion

Our study argues that temporal diversity in the fly optic lobe is a principled coding strategy tuned to the temporal statistics of natural motion. Circuits with high temporal diversity, typically combiming neurons with opposite polarities, achieve systematically higher coding performance for natural motion than circuits composed of more similar filters. Crucially, these benefits depend on stimulus statistics. Under white-noise inputs, the advantage of combined-polarity circuits disappears, and motion-circuit motifs with high diversity exhibit coding properties comparable to less diverse circuits (Figs. 4 and 5). In the motion pathway, these coding principles are reflected in specific anatomical motifs, including the columnar organization of medulla inputs onto direction selective dendrites. Together, our results link natural motion statistics to both temporal response diversity and the circuit architectures that implement it.

Visual systems are widely thought to be adapted to the ecological statistics of sensory inputs, and efficient-coding frameworks have been particularly influential in explaining responses to natural scenes [4,21,22,34,38–42]. Our work extends efficient-coding ideas to understand how visual systems handle natural motion. We leveraged on the temporal statistics of natural motion to provide a circuit-level normative account of how temporal diversity can improve coding under naturalistic, power-law input statistics.

Natural motion is inherently multiscale: luminance and contrast fluctuations distribute power across a broad range of temporal frequencies rather than around a single dominant timescale (Fig. 2). Under such scale-free input statistics, no single neuron can be simultaneously well matched to slow drifts and rapid transients. A more efficient strategy is therefore to distribute encoding across multiple temporal channels with complementary response properties and to combine them downstream. Fly visual circuits implement precisely such a strategy: integrating (single-polarity) and differentiating (double-polarity) filters span complementary temporal regimes (Fig. 1) and together support reliable reconstruction of natural inputs (Fig. 3). Although the temporal filters of medulla neurons can adapt with behavioral state [43], these changes appear to modulate response dynamics largely within polarity types. For example, neurons may become more transient without switching polarity type (from single to double polarity or vice versa). Multiscale inputs are therefore most effectively encoded not by a single neuron continuously adapting its own dynamics, but by pooling across neurons with heterogeneous temporal filters. In this regard, fly visual circuits resemble other biological systems in which several processes with distinct time constants collectively approximate scale-free dynamics over broad temporal ranges [24–27].

Temporal diversity as a coding strategy is likely relevant beyond the fly visual system. Diverse temporal receptive fields are a prominent feature of peripheral vision across species, including flies, mice, and zebrafish [4–7, 12, 13, 15, 17]. In the vertebrate retina, temporal receptive fields range from integrators to differentiators and vary across bipolar, amacrine, and ganglion cell populations. In flies, analogous diversity is already apparent in the lamina [8–10] and is further elaborated in the medulla [14, 15, 17, 18], where it has been particularly well characterized in motion-processing pathways [12]. Despite anatomical differences between insect and vertebrate visual systems, both appear to implement common computational strategies for the processing of visual information [44, 45]. This also includes the early decomposition of photoreceptor signals into parallel ON and OFF channels, that each contain cell types with distinct temporal filtering properties [34,46,47]. The specific anatomical realization of this strategy may differ across species. For instance, the number of temporally diverse pathways varies, with the mouse retina spanning around fifteen bipolar cells and about ∼ 30 functionally diverse ganglion cells [5,6], compared to the three temporally distinct lamina neurons or the eight temporally distinct medulla neurons in the fly. Still, the repeated emergence of diverse temporal filters early in visual processing suggests that cell type temporal diversity is a conserved design principle for matching neural representations to the multiscale structure of natural sensory inputs. In the vertebrate retina, efficient coding of spatiotemporal natural videos predicts the emergence of multiple cell types as the number of available retinal ganglion cells increased, with new types encoding progressively higher temporal frequencies [42]. Our results complement this picture in the mammalian visual system by showing that, given a repertoire of cell types, the specific combination of temporal filter polarities — rather than cell type number alone — determines circuit-level coding performance under natural motion statistics. Furthermore, recent work reports functional heterogeneity even within a single neuronal cell type, for example in zebrafish cones or within identified fly visual medulla cell types [36, 48–50], raising the possibility that temporal diversity can be generated at multiple organizational levels, from between to within cell type [48,50]. While the functional benefit of the latter is still unknown, our results show that diversity across identified cell types is already sufficient to yield measurable coding advantages at the circuit level. A key finding of our study is the tight connection between coding and anatomy of motion detection circuits. Temporally diverse filter combinations map naturally onto the spatial organization of medulla inputs to T4 and T5 dendrites (Fig. 5). This organization is stimulus dependent and becomes less evident under white-noise input statistics (Fig. 5), suggesting that the anatomical arrangement of presynaptic inputs onto direction-selective dendrites is not only a wiring solution for motion computation, but also an implementation of efficient temporal coding under natural motion statistics. At the same time, pathway-specific wiring cannot be derived from temporal statistics alone, and additional constraints (synaptic specificity, sign constraints, spatial pooling rules, and downstream nonlinearities) are likely required to recover the biologically realized subset of efficient solutions. Notably, natural images exhibit an over-representation of dark contrasts, making OFF contrasts more prominent than ON. Yet, the temporal structure of ON and OFF contrast fluctuation in natural motion appear largely indistinguishable [46], which may help explain why cross-pathway circuits can still appear efficient when evaluated for temporal coding alone.

Functional perturbation data provide independent support for this link between efficient coding and circuit organization. In particular, SNR predicted the effects of partial silencing of OFF-pathway medulla neurons on downstream LPTC responses [14], indicating that downstream motion signals are especially sensitive to perturbations that reduce presynaptic temporal diversity. In other words, the circuits that our framework identifies as best encoding natural motion are also those whose disruption has the strongest functional consequences. This agreement between efficient-coding predictions and perturbation data strengthens the link between temporal diversity at the medulla level and motion computations downstream.

Beyond functional predictions, circuits within and outside of motion detection pathway show different relationships between temporal diversity and synaptic co-organization. Only in motion-detection pathways circuits did high temporal diversity coincide with weak anti-correlations in synaptic counts across columns. One possible explanation is that these correlation structures reflect different anatomical and computational constraints. In circuitry presynaptic to T4/T5, temporally distinct inputs target different dendritic regions and contribute to the motion computations which require spatiotemporal comparisons between presynaptic inputs [37]. Here, synapse formation may involve local competition for resources such as dendritic territory or synaptic material, yielding weak anticorrelations between inputs that occupy adjacent but distinct compartments. Outside the motion pathway, where comparable compartmentalized spatiotemporal constraints may be absent or weaker, co-variation in synaptic strength may instead reflect coordinated recruitment of presynaptic channels that act together on shared downstream targets. This interpretation remains to be further tested, but it highlights that temporal diversity can be embedded in different circuit architectures depending on downstream computational demands.

Taken together, our findings support a view in which temporal diversity, circuit motifs, and natural stimulus statistics are tightly linked in the fly optic lobe. The early visual system appears to partition natural motion into complementary temporal channels whose joint readout improves coding efficiency, and this functional logic is reflected in the organization of motion-detection circuits and in the downstream effects of perturbing them. More generally, our results suggest that understanding why specific visual cell types exist, and why they are wired together in particular motifs, requires considering not only their intrinsic physiology or developmental origin, but also the temporal structure of the natural signals they evolved to encode.

## Acknowledgments

We thank Carlotta Martelli, Pradeepkumar Trimbake, Elizabeth Herbert and Claudia Cusseddu for comments on the manuscript, and are grateful to Shuai Shao, Kai Roth and Judith Parkinson-Schwarz for helpful discussions on the theoretical aspects of the work, and we thank the members of the Silies, Gjorgjieva, Schnaitmann, and Martelli labs for insightful discussions. The work was supported by funding from the Deutsche Forschungsgemeinschaft (DFG) via SPP2205 to MS and JG. This project has received funding from the European Research Council (ERC) under Horizon 2020 research and innovation programme (Grant agreement No. 101045003 ‘Adaptive Vision’)

## Author contributions

LR, JG, MS designed the project, LR and JG developed the theoretical methodology, LR performed research, analyzed and visualized data. LR and MS wrote the original draft of the manuscript, which was reviewed and edited by all authors.

## Methods

### Analytical description of naturalistic stimuli

The power spectral density of dynamic natural stimuli follows a power law in the temporal frequency domain, *∝* ω^*k*^, characterized by exponents ranging from k = −2 to k = − 1. We used this feature of natural motion to define our naturalistic stimuli. To have an analytically tractable problem, we used a Gaussian process described by the mixture in Fig. 2C, that is,

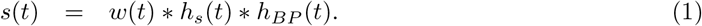

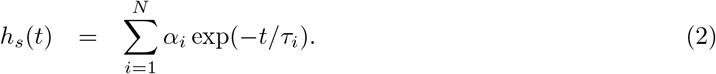

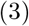

In Eq. 1, w(t) is a Gaussian white noise process, h_*s*_(t) is a low-pass filter mixture and h_*BP*_ (t) is the Butterworth band-pass filter, with magnitude

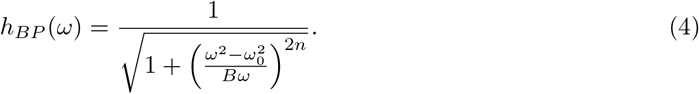

For the band-pass filter we set n = 4 with the parameters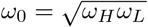 and B = ω_*H*_ − ω_*L*_. Signals from natural motion had most of the power within the frequency range of ω_*L*_ = 1Hz to ω_*H*_ = 100Hz which we then used as cut-off frequencies. With a mixture of N = 2 low-pass filters (Eq. 2) we fitted different power laws within this naturalistic frequency range (see Fig. 2).

### Neuron model

Lamina and medulla neurons exhibit graded responses that we modeled with a linear temporal filter, h_*i*_(t) and additive noise, *N* (0, σ_*N*_), i.e.,

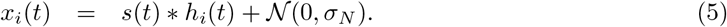

To fit the temporal filters, h_*i*_(t), of the fly optic lobe [12], we used the general biphasic function

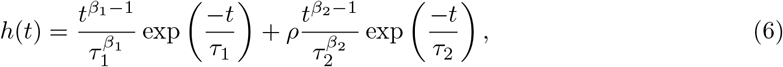

where the exponents β_*i*_ represent the order of the filters, and the parameters τ_*i*_ are the timescales of the two phases. The relative weight ρ is fit freely, including negative values. Fourth-order filters were used to fit temporal filters in the mammalian retina [30, 51]. Here we used a fourth order filter, β_1_ = 4, to account for the more transient initial response of neurons, and a third order filter, β_2_ = 3, to fit the slower phase following the initial peak (see Fig. S1). All filters were sign-aligned so that their dominant peak had positive polarity. The sign-aligned filter was then peak-normalized by dividing by its positive maximum, such that its maximal value was equal to 1.

The collective representation of all lamina and medulla filters lay in a two-dimensional feature space whose two main axes were the filter area and the half-life timescale. The total area, A, is the integral of the normalized filter, that is,

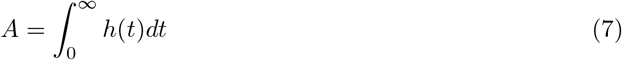

The half-life timescale, τ_*L*_ corresponds to the time at which the neuron filter integrates half of its total area, that is,

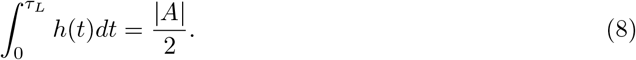

### Mean square error in networks of *N* neurons

For the analytical calculation below, we considered an unconstrained linear decoder in the Fourier domain. The model used for the natural-movie simulations is described separately in the corresponding subsection.

We optimized a linear decoder, F ^*\**^(t), to compute the estimated signal, s_est_(t), from a network of

N neurons, that is,

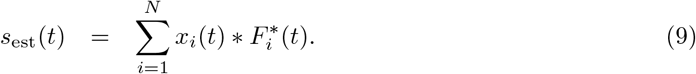

To measure the encoding performance of the network, we computed the mean square error (MSE) between the input and output signals,

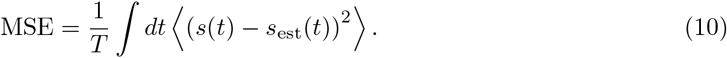

Expanding Eq. 10 yielded

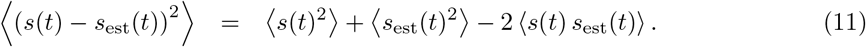

We defined the correlation functions

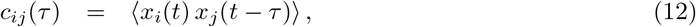

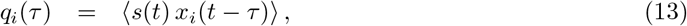

and denoted their Fourier transforms by C_*ij*_(ω) and Q_*i*_(ω), respectively. In the Fourier domain, the MSE can then be written as

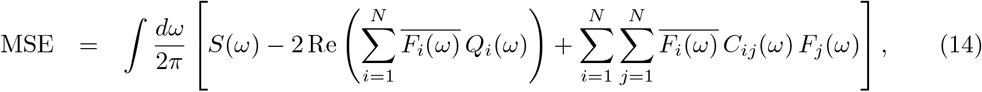

where S(ω) is the power spectral density of the stimulus, and F_*i*_(ω) is the Fourier transform of the optimal decoding filter 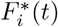.

We optimized the linear decoder by minimizing Eq. 14 with respect to 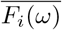, which yielded

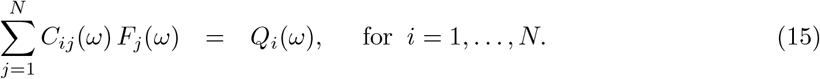

Thus, the optimal decoding filters satisfy the linear system

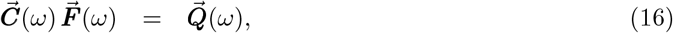

with

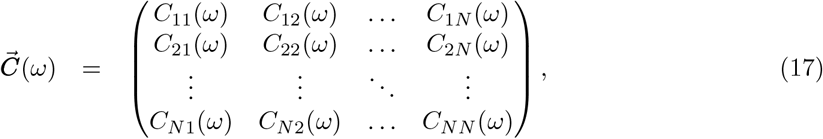

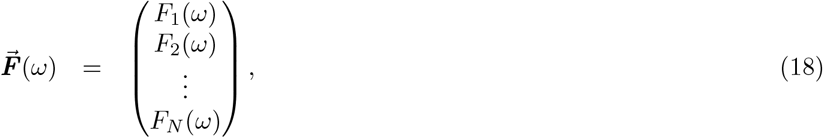

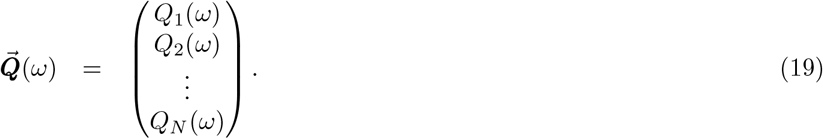

The solution of Eq. 15 was therefore

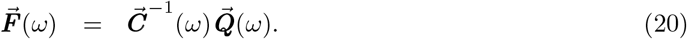

Equivalently, each optimal decoder could be written explicitly using Cramer’s rule as

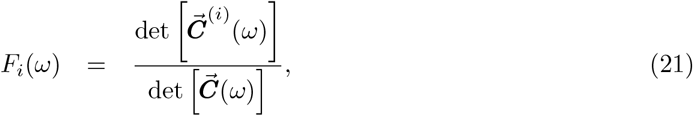

where 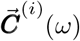 is the matrix obtained from 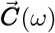 by replacing its i-th column by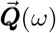. Finally, we defined the signal-to-noise ratio (SNR) as

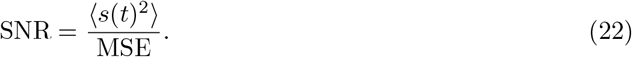

### Diversity coefficient

To quantify the diversity of temporal filters within circuits with N different neuronal types, we defined the diversity coefficient, D_*F*_, as a function of the difference of the areas and half-life timescales that described the filters in the feature space (see Fig. 2), for a circuit of two neurons that is,

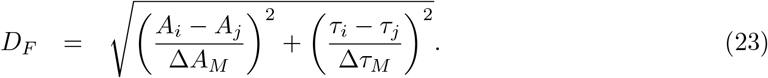

Each term of Eq. 23 was normalized and quantified the temporal diversity between pairs of neurons in networks of N different neuronal types. The parameters ΔA_*M*_ and Δτ_*M*_ corresponded to the maximum area and half-life timescale differences between the different neuronal types, such that the terms of Eq. 23 ranged from zero, e.g. same neurons, to unity.

### Generalized temporal filter space

To analyze coding strategies beyond the finite set of experimentally measured fly filters, we constructed a generalized ensemble of temporal filters in the two-dimensional feature space defined by filter area and half-life timescale. All filters were first sign-aligned such that the dominant peak was positive; if the largest-magnitude peak was negative, the full filter was multiplied by−1. Filters were then peak-normalized by their positive maximum. Under this convention, the area of the experimentally observed and generalized filter set considered here was restricted to the interval from 0 to 1, with values near 0 corresponding to strongly differentiating filters and values near 1 to strongly integrating filters. The half-life axis was restricted to the range of physiologically realistic timescales observed in fly data. We discretized this space on a grid whose resolution was chosen such that distinct experimentally measured fly filters remained separable at the grid level. For each grid point, we fitted the canonical biphasic filter function (Eq. 6) to reproduce the prescribed area and half-life timescale. This procedure also revealed a region of inadmissible solutions: filters with large area could not be realized with arbitrarily short half-life timescales, yielding the forbidden region shown in Fig. 4A. Generalized filters were classified as double- or single-polarity using a data-informed boundary at area A = 0.4, chosen to extend the experimentally observed classes approximately within one standard deviation of their respective distributions. This construction yielded an expanded filter ensemble spanning the fly-like temporal feature space beyond the measured cell types, comprising 97 double-polarity filters and 94 single-polarity filters. The similar number of filters in the two classes ensured that KDE- and KL-based comparisons were not dominated by strong class-size imbalance.

### Natural movie simulations

Natural-movie validation was performed using videos from the Chicago Motion Database [31], whose acquisition parameters and temporal statistics have been described previously [2, 31]. From each movie, we extracted one-dimensional temporal input traces by sampling image locations uniformly at random and taking the luminance value at that location across frames. Prior to extraction, frames were spatially smoothed with a Gaussian kernel (σ = 6 pixels), which served as a local spatial averaging step to suppress pixel-scale fluctuations. For each extracted trace, model responses were obtained by convolution with the corresponding neuronal temporal filters. The decoding filter was optimized separately for each video using the full stimulus trace and a causal lag window of 1 s. Reconstruction quality was quantified by the coefficient of determination, R^2^, between the decoded and original luminance trace.

### Connectome correlation analysis

We quantified correlations (Fig. 6E,J) in relative synaptic counts across columns of the same circuit motif. Absolute synaptic counts were normalized by the total number of synapses received by the postsynaptic neuron. For each presynaptic pair, we then measured the correlation between these relative synaptic counts across repeated instances of the same circuit in the fly eye.

Outside the motion detection pathway, common postsynaptic partners were selected using a fixed connectivity criterion. Specifically, a postsynaptic neuron was considered a common partner of two presynaptic neuron types only if it received more than 10% of its total synaptic input from each presynaptic neuron type, and if this connectivity motif was observed in at least 100 repeated instances across the fly eye. Presynaptic pairs that did not share any postsynaptic partner satisfying both criteria were excluded from the analysis.

## Supplementary Figures

**Figure S1:**
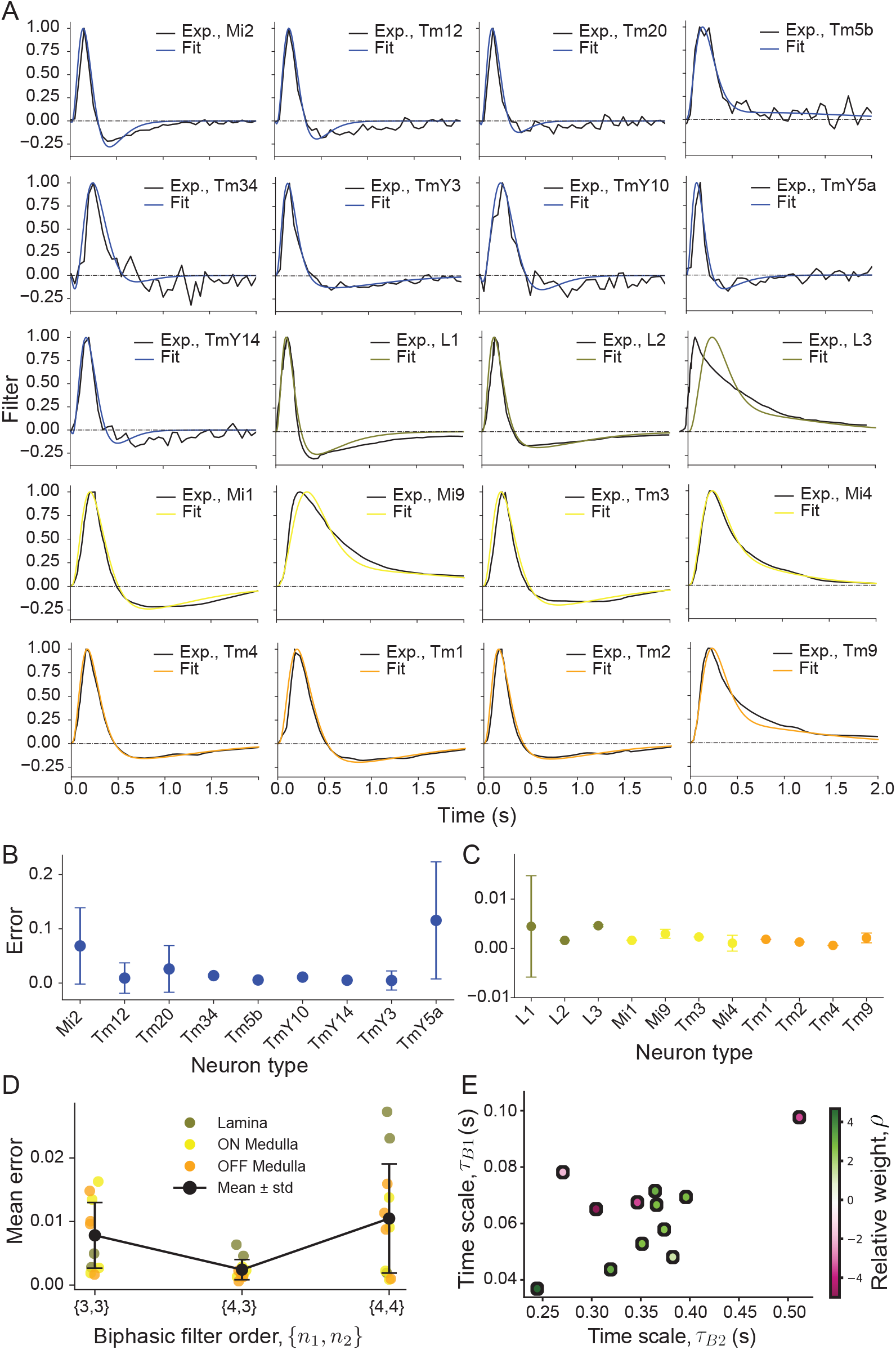
Supplementary figure on temporal filters in the fly visual system. **A** Temporal filters in the fly eye. Black curves correspond to data and colored curves show the fit function (see Eq. 6 and Fig. 1). Blue: medulla neurons outside the motion detection pathway; green/yellow/orange: Lamina/medulla-ON/medulla-OFF neurons within the motion detection pathway. **B**,**C** Error function for the fit of temporal filters outside the motion detection pathway (B) and within the motion detection pathway (C). **D** Mean error value for biphasic functions of different order in Eq. 6. Order n_1_ = 4 and n_2_ = 3 minimizes the fit error for lamina and medulla neurons within the motion detection pathway. **E** Parameter space of the filter fit function of Eq. 6. Each dot corresponds to the fit of the filters in Fig. 1C.

**Figure S2:**
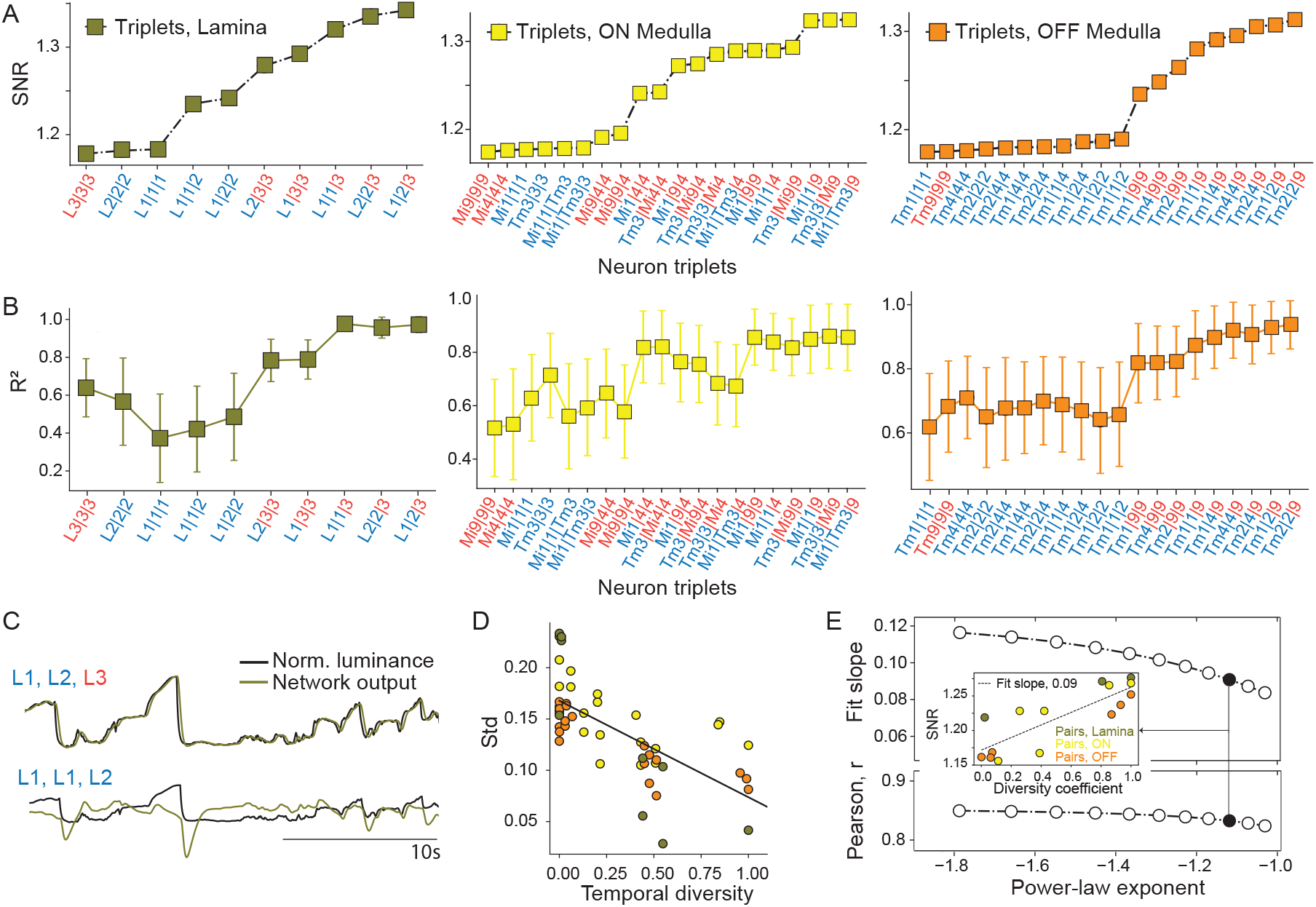
Encoding of naturalistic and natural inputs by triplets of lamina and medulla neurons. **A** Signal to noise ratio (SNR) of circuits of size N = 3 for all the combinations of lamina and medulla neurons in the ON and OFF pathway. Similar to Fig 3B,C. **B** Encoding performance for each lamina and medulla circuit, quantified by the R^2^ of a linear encoding model (temporal lag window of 1s). Networks are sorted as in A. Error bars extracted from simulations across 11 natural videos [2, 31]. **C** Luminance stimulus (black) and network response traces (green) for two network examples in the lamina with predicted high and low SNR in A. **D** Std of R^2^ versus temporal diversity coefficient across all networks (colors as in A). **E** Coding of lamina and medulla circuits across different power-law stimuli (see Fig. 2). For each stimulus, ranging from k = −1.8 to k = −1, we computed the linear fit between SNR and the diversity coefficient as in Fig. 3D. The upper plot shows the slope of the linear fit, and the lower plot shows the corresponding Pearson correlation. The inset shows an example of the SNR as a function of the temporal diversity coefficient for the power-law exponent marked in black.

